# Efficacies of sequenced monotherapies of *Mycobacterium avium* lung infection in mouse

**DOI:** 10.1101/2025.10.22.684001

**Authors:** Ruth A. Howe, Binayak Rimal, Jay Khandelwal, Chandra Panthi, Gyanu Lamichhane

## Abstract

**Background:** The incidence of non-tuberculous mycobacterial (NTM) infections has been rising and now exceeds tuberculosis in several countries. Among NTMs, *Mycobacterium avium* is the most common cause of chronic lung disease. Current guidelines recommend simultaneous administration of three or more antibiotics, modeled after tuberculosis treatment, but these regimens are limited by toxicity, poor adherence, and low cure rates. Importantly, unlike *M. tuberculosis, M. avium* is acquired from the environment rather than transmitted between humans, weakening the rationale for multidrug therapy as a strategy to suppress resistance at the population level.

**Methods:** To test an alternative treatment approach, we evaluated sequential monotherapy in a validated murine model of chronic *M. avium* lung infection. Mice were treated with either the standard triple-drug regimen of clarithromycin, ethambutol, and rifampicin or with sequential monotherapy: clarithromycin, bedaquiline, and clofazimine, with only one drug administered at a time for four-week intervals. Lung and spleen bacterial burdens were quantified, and minimum inhibitory concentrations (MICs) were determined for isolates recovered during treatment to assess resistance emergence.

**Results:** Sequential monotherapy achieved reductions in lung bacterial burden equivalent to those of the standard multidrug regimen and prevented extrapulmonary dissemination. Notably, no increase in MICs was observed for clarithromycin, bedaquiline, or clofazimine across treatment phases, indicating that sequential monotherapy did not select for resistant clones.

**Conclusions:** These findings provide the first experimental evidence that sequential monotherapy can deliver efficacy comparable to multidrug therapy for *M. avium* disease without promoting resistance. This proof-of-concept supports further investigation of sequencing strategies as a potentially more tolerable alternative to current regimens.

## INTRODUCTION

The incidence of non-tuberculous mycobacterial diseases is rising globally and recently overtook tuberculosis in the US [1,2]. *Mycobacterium avium* complex (MAC) is the most isolated non-tuberculous mycobacterial pathogen in humans [3,4]. MAC is comprised of closely related species with *M. avium, M. intracellulare* and *M. chimaera* as human pathogens. Among MAC, *M. avium* is most commonly isolated from patients and *M. chimaera* is rare [3]. MAC are slow-growing organisms requiring growth conditions similar to *M. tuberculosis* in laboratory. The ability of MAC to thrive in urban water supply systems has been considered one of the contributing factors to the increasing incidence of their infections globally [5].

Infections with *M. avium* primarily manifest as chronic lung disease with symptoms that include chronic cough, fatigue, weight loss and reduced lung function resembling—and often initially misdiagnosed as—tuberculosis. Individuals with structural lung conditions such as cystic fibrosis, bronchiectasis and chronic obstructive pulmonary disease are at an elevated risk of contracting *M. avium* infection. In immunocompromised patients, particularly those with advanced HIV/AIDS, *M. avium* can cause disseminated disease with fever, anemia and hepatosplenomegaly [3,6]. *M. avium* infection significantly impairs quality of life, can lead to progressive lung damage, and increases morbidity and mortality. A 2018 systematic review of 14 studies (9,035 patients) reported a 5-year all-cause mortality rate of 27% (95% CI 21-38%) [7]. A 2020 study in Japan (1,445 patients) found 5- and 10-year mortality rates of 24% and 46%, respectively [8]. One of the primary reasons for poor prognosis of *M. avium* diagnosis is the lack of effective treatment. Liposomal formulation of amikacin is the only FDA-approved drug for treating *M. avium* disease.

The current treatment guidelines for *M. avium* lung disease rely heavily on expert opinion shaped by clinical experience. Experts who developed these guidelines have noted disagreements over treatment regimens due to varying individual experiences [9]. Current guidelines recommend combination therapy comprising of three or more drugs, typically a macrolide as the backbone drug (such as azithromycin or clarithromycin), ethambutol, and rifampicin (± aminoglycoside), administered for at least 12 months after sputum culture conversion to negative [6,10]. However, toxicities, poor adherence and high relapse rates contribute to an average cure rate of only 31% and refractory disease is common, necessitating lifelong patient follow-up [3,11].

Treatment related adverse events occur in 60-70% of patients and 30-70% of patients discontinue at least one drug from their initial regimen due to inability to tolerate [12]. In a study of 364 MAC-treated patients, long-term administration of two or more drugs led to hepatotoxicity in ∼19.5% (median onset ∼55 days), leukopenia (∼20% at ∼41 days), thrombocytopenia (∼28.6% at ∼61 days), ocular toxicity (∼7.7% at ∼278 days), and renal impairment (∼12.4% at ∼430 days) [13]. Most regimens include three or more antibiotics, each with well-documented toxicities, ranging from hepatotoxicity and cytopenias to ototoxicity, optic neuritis, neuropathy and gastrointestinal upset [3,14]. Drug-drug interactions, especially involving rifamycins reducing macrolide levels, further complicate pharmacokinetics and contribute to adverse event exacerbation. Collectively there is significant evidence linking long-term intake of multiple drugs simultaneously to the cumulative toxicity and intolerability of the currently recommended therapy [12].

The currently recommended treatment for *M. avium* disease involves a multidrug regimen during both the intensive and continuation phases, modeled after the treatment strategy for tuberculosis [10]. In tuberculosis, the use of multiple drugs is essential to prevent the emergence of drug-resistant strains, which pose a significant public health threat due to their potential for human-to-human transmission. However, this public health rationale does not apply to *M. avium* disease. Unlike *M. tuberculosis, M. avium* is not transmitted between humans; instead, it is acquired from environmental sources [5,15]. As a result, any drug-resistant *M. avium* that emerges during treatment remains confined to the individual patient and does not pose a broader public health risk. While drug resistance has been shown to emerge in *M. avium* in the course of prolonged macrolide monotherapy or interrupted multi-agent regimens in complex disease, there is less evidence suggesting emergence of resistance in non-disseminated, noncavitary disease [3,6,16,17]. Therefore, the justification for using multidrug therapy to prevent the spread of resistance, a cornerstone of tuberculosis control, is far less compelling in the case of *M. avium*, where humans represent a dead-end host in the pathogen’s evolutionary cycle. While it is theoretically possible that drug-resistant *M. avium* shed by an infected individual could enter environmental reservoirs—such as water systems or soil—and later be acquired by another person, this route of transmission has not been clearly demonstrated. Thus, while environmental persistence and potential transmission cannot be completely ruled out, this pathway remains speculative. An additional rationale for using multidrug therapy is the potential for synergism among drugs, which can accelerate *M. avium* clearance and enhance overall efficacy compared to monotherapy.

We hypothesize that administering drugs sequentially as monotherapy may mitigate the intolerability associated with the simultaneous use of multiple antibiotics. To test this, we compared the efficacy of the standard multidrug regimen—comprising clarithromycin, ethambutol, and rifampicin—with a sequential monotherapy approach. In one version of this approach, treatment began with clarithromycin monotherapy, followed by bedaquiline, and concluded with clofazimine. In the second version, the sequence was reversed, starting with clofazimine, followed by bedaquiline, and ending with clarithromycin. Sequenced monotherapy with clarithromycin, ethambutol, and rifampicin would provide a direct comparison to the standard-of-care regimen. However, previous studies have shown that rifampicin is ineffective for treating *M. avium* disease [18–20], and ethambutol primarily serves to minimize emergence of macrolide resistance [16] making them less suitable for monotherapy evaluation compared to bedaquiline and clofazimine, which demonstrate strong standalone activity [21] and low rates of resistance emergence in patients [22]. To assess the potential for resistance emergence, we measured the minimum inhibitory concentrations (MIC) of the administered drugs against *M. avium* progenitors recovered after treatment. All studies were conducted using an established mouse model of *M. avium* lung infection that is validated for evaluating treatment efficacy [23,24].

## MATERIALS AND METHODS

### *M. avium* strain, growth media and conditions

The *M. avium* strain ATCC 700898, also known as MAC101, and widely used a reference of *M. avium* in laboratory studies [25–27] was used in this study. This strain was purchased from American Type Culture Collection (Manassas, Virginia) in 2022, and the first passage stock, maintained in -80 °C, was used for all experiments. MAC101 and its progenitor colonies recovered from mouse lungs and spleens were cultured in Middlebrook 7H9 broth (Difco, catalog no. 271310) supplemented with 0.5% glycerol, 0.05% Tween-80, and 10% oleic acid–albumin–dextrose–catalase (OADC; BD, catalog no. 212351). Cultures were incubated at 37°C with shaking at 220 RPM, according to established protocols [28]. For recovery of MAC101 and its progenitors from mouse tissues, 10-fold serial dilutions of lung or spleen homogenates were inoculated onto Middlebrook 7H11 selective agar (Difco, catalog no. 283810) supplemented with 0.5% glycerol, 0.05% Tween-80, 10% OADC, and 50 μg/mL cycloheximide (Sigma-Aldrich, catalog no. C7698). Plates were incubated at 37°C for 10–14 days, in accordance with Clinical and Laboratory Standards Institute (CLSI) guidelines [29].

### Infection and treatment efficacy assessment in mice

A validated mouse model of chronic *M. avium* lung infection, as described by Andrejak et al. and subsequent studies [21,23,24], was used to evaluate treatment efficacy. Female BALB/c mice (n = 70, 5–6 weeks old) were obtained from Charles River Laboratories (Wilmington, Massachusetts) and acclimatized for 7–10 days in a biosafety level 2 vivarium prior to infection. To establish infection, MAC101 culture in exponential growth phase (A_600nm_ = 1.00–1.60) was diluted in Middlebrook 7H9 broth to A_600nm_ = 1.0. Ten milliliters of this suspension were aerosolized into the chamber of a Glas-Col Inhalation Exposure System A4212 (Glas-Col, Terre Haute, Indiana) using a nebulizer. The exposure sequence included 15 minutes of pre-heat, 30 minutes of aerosolization, 30 minutes of aerosol decay, and 15 minutes of ultraviolet surface decontamination. Mice were infected simultaneously through natural respiration of the aerosolized suspension. Following infection, mice were randomized into groups of five per cage.

*M. avium* implantation load in the lungs was determined by sacrificing five mice one day post infection (designated ‘week -4’). Lungs were aseptically extracted, homogenized in 1× PBS with 2 mm glass beads, and disrupted by bead-beating for 30 seconds at 4,000 RPM (Minilys, Bertin Instruments). Serial dilutions (0.1 mL) were inoculated onto standard petri dish containing Middlebrook 7H11 selective agar, incubated at 37°C for 14 days, and colony-forming units (CFUs) enumerated. A second set of five mice was sacrificed four weeks later (“week 0”), marking the initiation of antibiotic treatment. Thereafter, lung bacterial burden was determined at weeks +4, +8, and +12, with five mice evaluated per treatment group per timepoint.

### Treatment Groups

Mice were assigned to one of four treatment groups (n = 15 per group; Figure 1). All drugs and controls were administered by oral gavage in 200 μL volumes using a 22-gauge curved gavage needle (2-mm tip; Gavageneedle.com, AFN2425C) attached to a 1-mL slip-tip syringe (Becton & Dickinson, catalog no. 309659). Treatments were given once daily, seven days per week, for up to 12 weeks.

**Figure 1.**
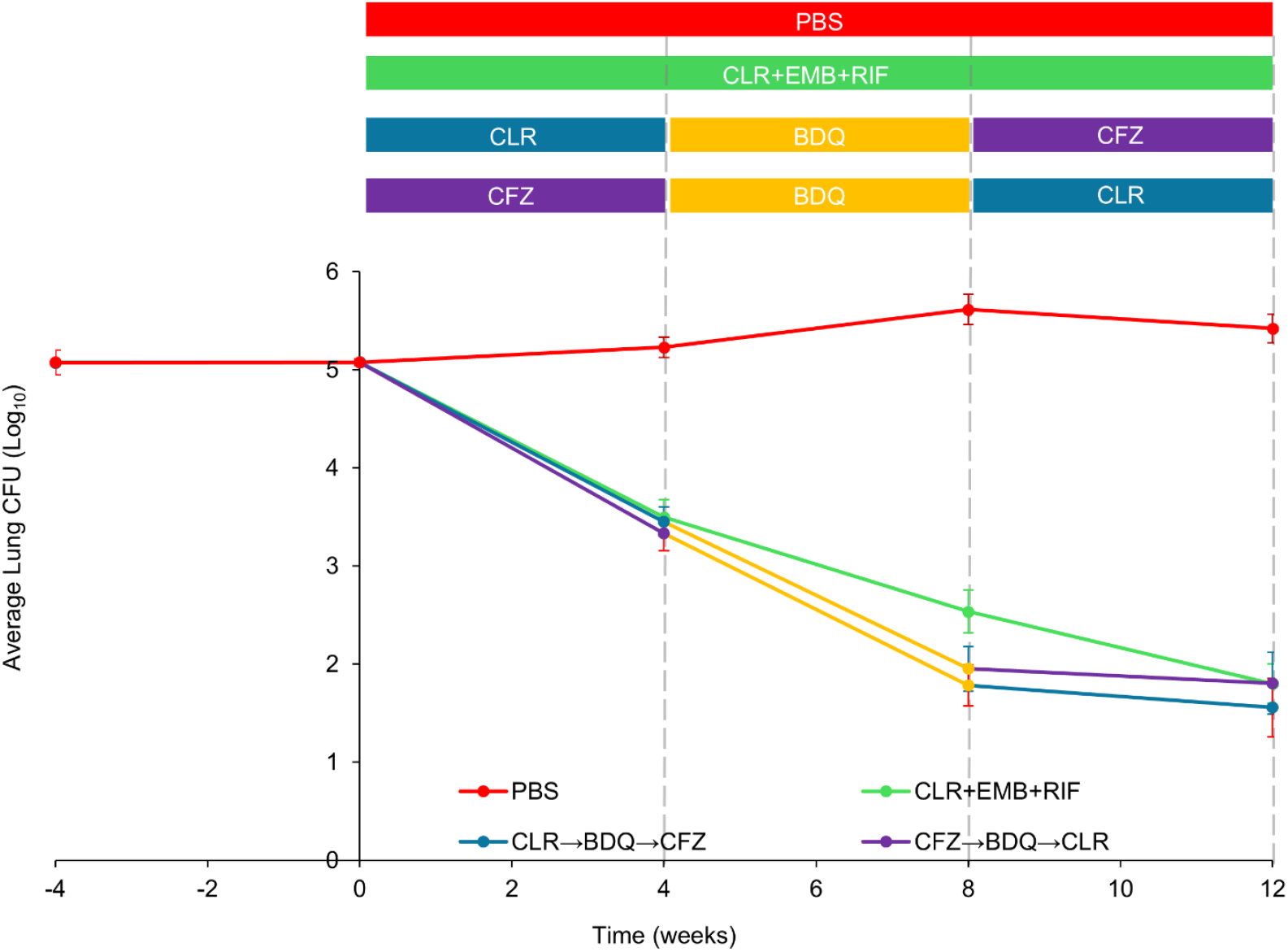
Evaluation of sequential monotherapies versus standard-of-care in a mouse model of *M. avium* lung infection. The top panel depicts the study design. The first group received phosphate-buffered saline (PBS) as control. The second group received the standard-of-care regimen: clarithromycin (CLR, 100 mg/kg), ethambutol (EMB, 100 mg/kg), and rifampicin (RIF, 20 mg/kg) administered daily. The third group received sequential monotherapy: CLR (100 mg/kg) for 4 weeks, followed by bedaquiline (BDQ, 25 mg/kg) for 4 weeks, and clofazimine (CFZ, 25 mg/kg) for 4 weeks. The fourth group received the same agents and doses in reverse order (CFZ → BDQ → CLR). Each treatment group included 5 mice per timepoint.

Mice in the first group, designated no treatment control, received 200 μL bolus 1xPBS, once daily via oral gavage as 1xPBS was used as solvent for antibiotics used in this study. Mice in the second group, representing standard-of-care, received once daily clarithromycin (100 mg/kg), ethambutol (100 mg/kg), and rifampicin (20 mg/kg) for 12 weeks, with drug doses established in a prior study approximating their exposures in humans [23]. Mice in the third group, representing an experimental sequenced monotherapy group received clarithromycin (100 mg/kg) once daily during the first four weeks, followed by bedaquiline (25 mg/kg) for four weeks, and finally clofazimine (25 mg/kg) for four weeks. Mice in the fourth group, also representing experimental sequenced monotherapy group received the same drugs, doses, and dosing frequency, but administered in the reverse order: clofazimine first, then bedaquiline, and concluding with clarithromycin.

### Ethics statement

All animal procedures complied with national guidelines and were approved by the Johns Hopkins University Animal Care and Use Committee (protocol MO23M163).

### Data analysis

Raw lung CFU data were analyzed, and the mean ± standard deviation was calculated for each treatment group at each timepoint. These results were graphed as dot plots. Variance between groups was analyzed by one-way ANOVA with multiple comparisons (Supplementary Table 1). A p-value ≤ 0.05 was considered statistically significant. A non-inferiority test with a margin of 15% was used to compare the experimental treatments against the standard-of-care at the final timepoint using net CFU reduction the groups (Supplementary Table 2).

### Antibiotics Preparations

All antibiotics were prepared under sterile conditions. For Clarithromycin (Sigma-Aldrich, catalog no. C9742), the amounts of the powdered form necessary for administration to mice for each day were weighed into 5 ml polypropylene tubes prior to treatment initiation and stored at 4°C. Each day, the daily aliquot was retrieved, mixed with 0.05% agarose at 4°C to prepare a concentration of 10 mg/mL, and vortexed to generate a homogeneous suspension that appears cloudy white. An 0.05% agarose solution was prepared by adding 50 mg Bacto agar (BD, catalog no. 214010) to 100 mL 1x phosphate buffered saline (PBS), pH 7.4 (Quality Biologicals, catalog no. 114-058-101), autoclaving for 10 min at 121°C and stored at 4°C until use.

Similarly, for ethambutol (Sigma-Aldrich, catalog no. E4630), amounts necessary for daily administration were weighed into 5 ml polypropylene tubes and stored at 4°C. Each day, the daily aliquot was retrieved, and 10 mg/mL solution was prepared in 1xPBS. This preparation appears colorless and clear.

For rifampicin (Sigma-Aldrich, catalog no. R7382), the amounts of powder necessary for each week were weighed into 50 ml polypropylene tubes and stored at 4°C. At the beginning of each week, the weekly aliquot was retrieved, mixed with 0.05% agarose at 4°C to prepare a suspension at 2 mg/mL, and vortexed for 5 minutes. This preparation appears as dark red homogeneous suspension. The aliquot necessary for each day was transferred to 5 ml tubes and stored at 4°C until use.

For bedaquiline, powdered form bedaquiline fumarate (CAS no. 845533-86-0, Octagon Chemicals Ltd) was used. The amounts of the powder necessary for each week were weighed into a 100 ml borosilicate bottle, the precise volume of 20% 2-hydroxypropyl-β-cyclodextrin solution was added and dissolved by stirring with a magnetic stirrer for three hours at 4°C to prepare 2.5 mg/mL solution which appears transparent. Aliquots necessary for each day were transferred to 5 ml tubes and stored at 4°C until use. A 20% 2-hydroxypropyl-β-cyclodextrin (HPCD) (Sigma-Aldrich, catalog no. 332593) solution was prepared as described [30]. Briefly, 20 g of HPCD powder was transferred to a 100-mL borosilicate bottle, and 75 mL of sterile deionized water was added and stirred with a magnetic stirrer until a clear solution was obtained (∼30 min). Approximately 1.5-mL of 1 N HCl was added to bring pH to 2.0, and the final volume was brought to 100 mL by adding sterile DI water. This solution was filtered through a 0.22-mm acetate cellulose filter and stored at 4°C until use.

For clofazimine (Sigma-Aldrich, catalog no. C8895), the weekly amount of powder was weighed into 50 mL polypropylene tubes and stored at 4°C. At the beginning of each week, the weekly aliquot was retrieved, mixed with 0.05% agarose at 4°C to prepare a concentration of 2.5 mg/mL, and vortexed for 5 minutes at maximum speed. This suspension was then sonicated at 50% power for 15 seconds per cycle, with 2-3 cycles, until a matte red, opaque, homogeneous colloidal suspension was achieved. Aliquots necessary for each day were transferred to 5 ml tubes and stored at 4°C until use.

### Determination of MICs

Minimum inhibitory concentrations (MICs) for clarithromycin, ethambutol, rifampicin, bedaquiline, and clofazimine were determined using broth microdilution according to CLSI guidelines for slow-growing non-tuberculous mycobacteria [29]. A 10 mg/mL stock solution of each drug was prepared by dissolving the drug in dimethyl sulfoxide and diluted in Middlebrook 7H9 broth supplemented with 0.5% glycerol and 5% oleic acid-albumin-dextrose-catalase enrichment (without Tween-80) to make working solutions of 512 μg/mL. This working solution was serially 2-fold diluted in the same Middlebrook 7H9 broth in 96-well microtiter plate to generate final drug concentrations ranging from 128 μg/mL to 0.125 μg/mL.

At each timepoint (weeks 0, +4, +8, +12), two progenitor colonies of MAC101 were recovered from each mouse from each timepoint and treatment group from Middlebrook 7H11 agar plates used for CFU enumeration. Therefore, a total of 130 distinct CFUs were recovered representing two per mouse from 65 mice. These colonies were grown in Middlebrook 7H9 broth to prepare stocks and archived in -80 °C. The stocks were used to inoculate Middlebrook 7H9 broth and each MAC101 progenitor was grown to exponential phase, 10^5^ CFU was inoculated into each well of a 96-well microtiter plate with a final volume of 150 μL per well, alongside negative (broth only) and positive (broth with parent MAC101) controls, and incubated at 37°C for 10 days without shaking, following CLSI guidelines. MICs were determined using the Sensititre Manual Viewbox as the lowest concentration preventing visible growth. Assessment with all five drugs were performed simultaneously on the same plate for each strain. Therefore, 130 96-well microtiter plates were used to complete this study.

## RESULTS

### Efficacy of Sequenced Monotherapy

Mice were infected with aerosolized *M. avium* MAC101 to mimic the natural respiratory route of infection in humans, following the model described previously [23]. The scheme of the study is outlined in figure 1. Treatment began at the start of the fifth week post-infection. Two comparator arms were included. The first, representing the standard-of-care regimen, received once-daily doses of clarithromycin (100 mg/kg), ethambutol (100 mg/kg), and rifampicin (20 mg/kg) for 12 weeks, with dosing designed to approximate human drug exposures as established in an earlier study [23]. The second comparator arm received only phosphate-buffered saline (PBS)—the solvent for most drugs in this study—serving as the no-treatment control. Two experimental arms tested sequential monotherapy regimens, with each drug administered alone for four weeks before switching to the next. In the first experimental group, mice received clarithromycin monotherapy (100 mg/kg) for four weeks, followed by bedaquiline monotherapy (25 mg/kg) for four weeks, and finally clofazimine monotherapy (25 mg/kg) for four weeks. The second experimental group received the same drugs, doses, and dosing frequency, but in the reverse order: clofazimine monotherapy first, then bedaquiline monotherapy, and finally clarithromycin monotherapy. Lung MAC101 burden in each treatment group is shown in figure 1 (and statistical comparisons between groups are included in Supplementary Table 1).

At the time of infection, the mean lung MAC101 burden was 5.07 log_10_ CFU (Figure 1). This infection was allowed to progress for four weeks to establish a chronic infection. In the PBS-only control group, the lung bacterial burden remained stable throughout the study, confirming persistence of chronic infection as in prior studies [21,23]. In the standard-of-care group, the mean MAC101 lung burden declined steadily over the 12-week treatment period, resulting in a net reduction of 3.27 log_10_ CFU. In the first experimental group, clarithromycin monotherapy during the initial four weeks reduced the lung burden by 1.62 log_10_ CFU—similar to the 1.58 log_10_ CFU reduction seen in the standard-of-care group over the same period. During the second phase, bedaquiline monotherapy produced an additional 1.50 log_10_ CFU reduction, compared with 0.96 log_10_ CFU in the standard-of-care group. In the final phase, clofazimine monotherapy yielded a further 0.15 log_10_ CFU reduction, leading to a total reduction of 3.27 log_10_ CFU—identical to the net effect of the standard-of-care regimen. A non-inferiority assessment demonstrated that CFU reduction by this sequenced monotherapy was non-inferior to the standard-of-care (Supplementary Table 2).

In the second experimental group, clofazimine monotherapy in the first four weeks reduced the bacterial burden by 1.75 log_10_ CFU, again comparable to the 1.58 log_10_ CFU reduction with standard-of-care. Bedaquiline monotherapy in the second phase yielded an additional 1.55 log_10_ CFU reduction, and clarithromycin monotherapy in the final phase produced a further 0.23 log_10_ CFU reduction. This regimen achieved a total reduction of 3.52 log_10_ CFU over the study period—matching the efficacy of the standard-of-care triple-drug regimen with an overall efficacy being non-inferior to the standard-of-care.

In addition to measuring MAC101 burden in the lungs, we also quantified its burden in the spleen to evaluate the impact of each treatment on extrapulmonary dissemination. At the start of treatment, the mean spleen burden was 1.88 log_10_ CFU. In mice receiving PBS only, the spleen burden increased steadily during the treatment period, reaching 3.97 log_10_ CFU. In contrast, in all other treatment groups, no CFUs were detectable after the first four weeks of treatment and they remained undetectable thereafter.

### Assessment of Drug-Resistance Emergence

Monotherapy for *Mycobacterial* infections is generally expected to promote the emergence of drug resistance, ultimately leading to refractory disease. This expectation underpins the current clinical recommendation for simultaneous administration of multiple drugs not just in *M. tuberculosis* and *M. abscessus*, but for all subtypes of MAC. Based on this rationale, we hypothesized that MAC101 colonies recovered at the end of each phase of monotherapy in our experimental groups would show a significant increase in MICs for the drugs to which they were exposed, indicating resistance emergence.

To test this hypothesis, we determined the MICs of clarithromycin, ethambutol, rifampicin, bedaquiline, and clofazimine against MAC101 progenitors recovered from mouse lungs after 4, 8, and 12 weeks of treatment. Ten independent colonies (two per mouse) were tested at each timepoint per treatment group, yielding 130 colonies from 65 mice. At baseline (4 weeks of infection, prior to drug exposure), the median MICs were: clarithromycin 1 μg/mL, ethambutol 16 μg/mL, rifampicin 1 μg/mL, bedaquiline 0.125 μg/mL, and clofazimine 1 μg/mL (Figure 2). In the standard-of-care group (clarithromycin, ethambutol, rifampicin daily for 12 weeks), median MICs for clarithromycin, ethambutol, bedaquiline, and clofazimine remained unchanged, while rifampicin increased slightly from 1 to 2 μg/mL. Thus, no significant loss of susceptibility occurred for these agents.

**Figure 2.**
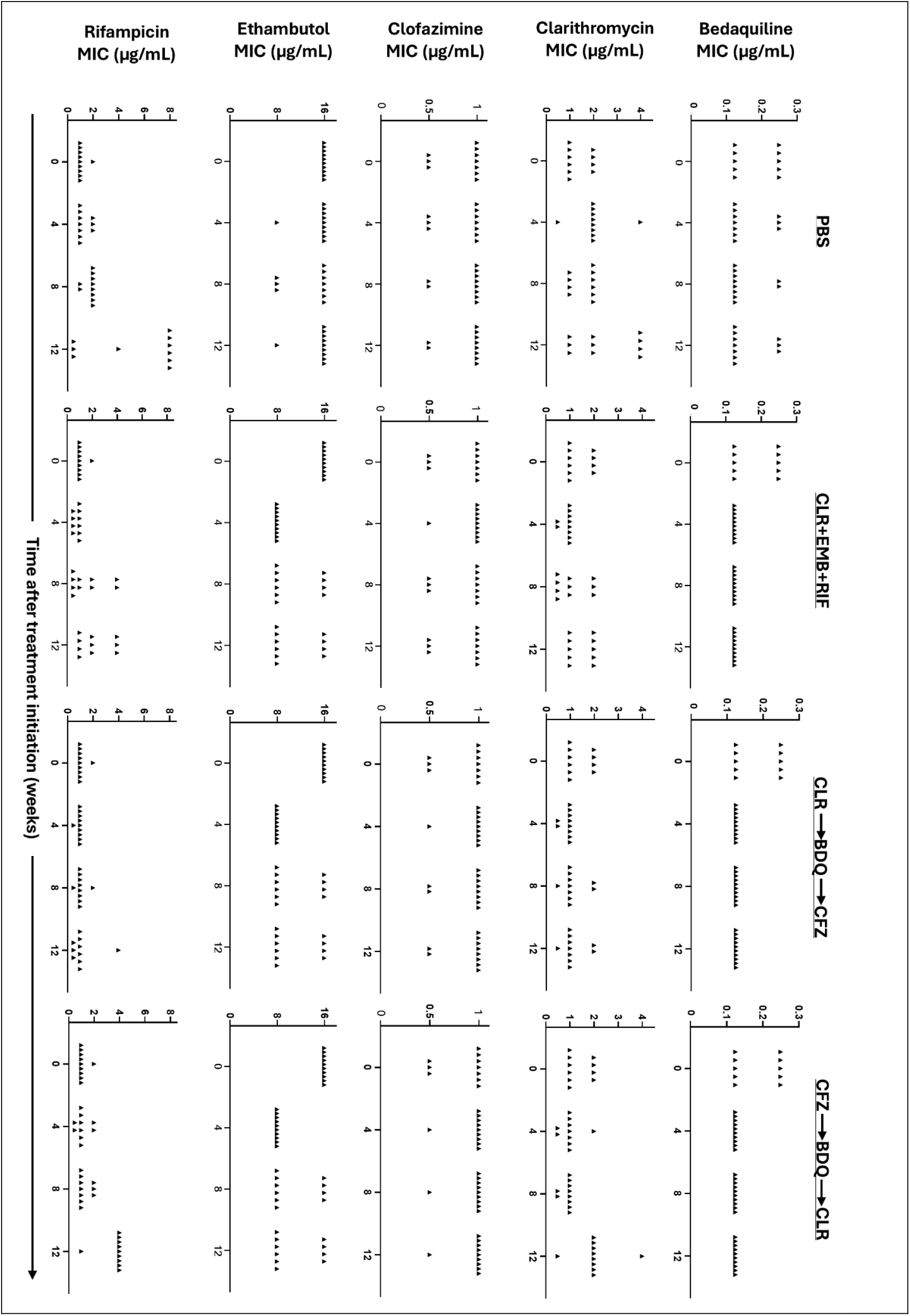
Minimum inhibitory concentrations (MICs) of antibiotics against *M. avium* progenitors recovered from mouse lungs. MICs of bedaquiline (BDQ), clarithromycin (CLR), clofazimine (CFZ), ethambutol (EMB), and rifampicin (RIF) were determined for progenitors of *M. avium* strain MAC101 isolated from each treatment group across all time points (n = 10 progenitors per time point per treatment group; n = 5 mice per time point per treatment group). The ‘PBS’ group received phosphate-buffered saline and therefore served as the no-treatment control. The ‘CLR+EMB+RIF’ group received a triple-drug regimen of clarithromycin, ethambutol, and rifampicin. The ‘CLR→BDQ→CFZ’ group received sequential monotherapy consisting of clarithromycin for the first four weeks, followed by bedaquiline for the next four weeks, and clofazimine for the final four weeks. The ‘CFZ→BDQ→CLR’ group received the same three drugs as sequential monotherapy but in reverse order.

### First experimental group – clarithromycin → bedaquiline → clofazimine sequence

After four weeks of clarithromycin monotherapy, MICs for all drugs—including clarithromycin—remained unchanged, directly contradicting the hypothesis that clarithromycin monotherapy would select resistant clones. The same was true following four weeks of bedaquiline monotherapy, providing strong evidence that short-term bedaquiline monotherapy does not induce resistance. After the final phase of clofazimine monotherapy, MICs again remained unchanged for all drugs except ethambutol, whose median MIC decreased from 16 to 8 μg/mL. Notably, clofazimine resistance did not emerge. Overall, sequential monotherapy with clarithromycin, bedaquiline, and clofazimine for four weeks each did not result in resistance to any of these drugs.

### Second experimental group – clofazimine → bedaquiline → clarithromycin sequence

Clofazimine monotherapy for four weeks left MICs for all drugs unchanged, ruling out resistance emergence. The same was observed after four weeks of bedaquiline monotherapy. Following the final phase of clarithromycin monotherapy, median MIC for clarithromycin rose from 1 to 2 μg/mL, but this increase was too small to indicate resistance, as defined by the Clinical and Laboratory Standards Institute breakpoints [29]. Indeed, among the 10 colonies recovered at this point, clarithromycin MICs ranged from 0.5 to 4 μg/mL, with only one colony reaching 4 μg/mL (Figure 2). Ethambutol and rifampicin MICs also shifted (16 → 8 μg/mL and 1 → 4 μg/mL, respectively) despite these drugs not being administered during this phase, suggesting changes unrelated to direct selection pressure. In summary, across both experimental regimens, four-week monotherapy with clarithromycin, bedaquiline, or clofazimine did not lead to the emergence of resistance to these drugs in *M. avium* MAC101. These findings challenge the assumption that short-term monotherapy inevitably selects for drug-resistant clones in this model.

### Gross body weights of mice

In severe mycobacterial infections, progressive bacterial proliferation and the associated chronic inflammatory response typically lead to gradual weight loss, though weight may be stable in early or mild disease [31,32]. Consequently, declining body weight correlates with disease exacerbation, whereas stabilization or recovery of weight can reflect therapeutic benefit and improved systemic health. Gross body weight can also serve as an indicator of treatment tolerability, as loss of body weight is one of the most readily observable clinical signs when antibiotics are not well tolerated, thought to be predominantly due to effects on the microbiome and nutrient uptake [33,34]. Accordingly, whole-body weight serves as a convenient, non-invasive, and broadly informative marker of both mycobacterial disease progression and treatment tolerability in murine models of mycobacterial infection.

We monitored gross body weight throughout the study to evaluate the effect of each treatment. This data is shown in figure 3 (and statistical assessments of the mean body weights between treatment groups are included in Supplementary Table 3). Consistent with mild chronic pulmonary *M. avium* infection, untreated animals had slow, stable weight gain over the course of the study. However, by the end of the first two weeks, mean body weight had decreased by 5% in mice treated with the standard-of-care regimen (clarithromycin, ethambutol, and rifampicin) but had increased by 8% in mice treated with clarithromycin monotherapy. This striking difference occurred despite both groups receiving the same dose of clarithromycin, suggesting that the addition of two other drugs in the standard-of-care regimen was associated with an early reduction in body weight. Although mice in the standard-of-care group gradually regained weight and matched the clarithromycin group during the middle portion of treatment, by study completion, mice that began with clarithromycin monotherapy had a higher mean body weight than those in either the standard-of-care group or the group receiving clofazimine monotherapy at the outset. Mean body weight also increased steadily in mice that received no antibiotics or monotherapies consisting of clofazimine followed by bedaquiline and clarithromycin.

**Figure 3.**
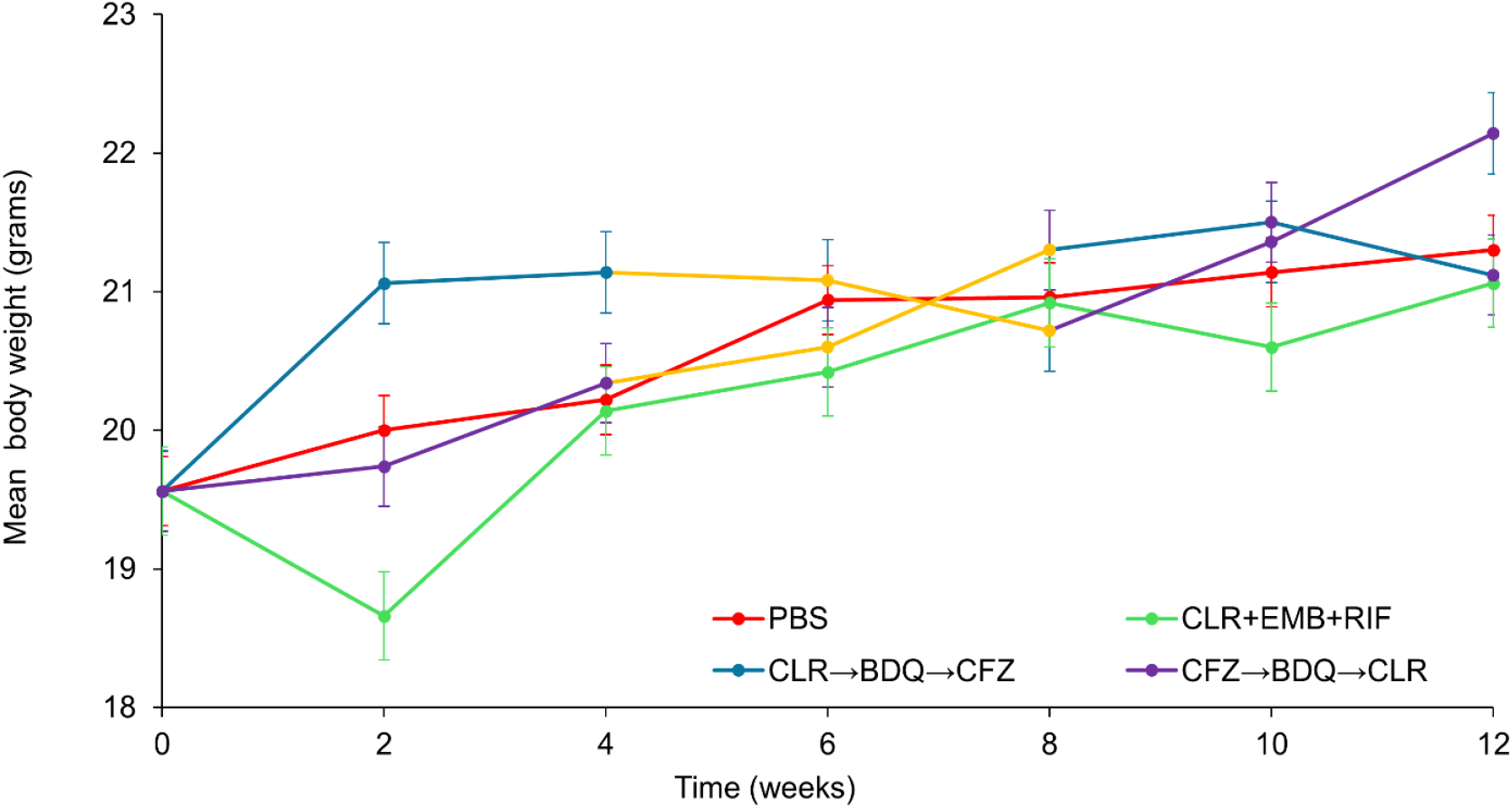
Body weight trajectories of *M. avium*–infected mice under different treatment regimens. Mean body weights ± standard error are shown at two-week intervals (n = 5 mice per time point per treatment group). The ‘PBS’ group (red) received phosphate-buffered saline and served as the no-treatment control. The ‘CLR+EMB+RIF’ group (green) received a triple-drug regimen of clarithromycin, ethambutol, and rifampicin. The ‘CLR→BDQ→CFZ’ group received sequential monotherapy consisting of clarithromycin (blue) for the first four weeks, followed by bedaquiline (orange) for the next four weeks, and clofazimine (purple) for the final four weeks. The ‘CFZ→BDQ→CLR’ group received the same three drugs as sequential monotherapy but in reverse order.

Overall, body weight trends revealed that treatment initiation with clarithromycin monotherapy was associated with the most pronounced early weight gain, whereas the addition of other drugs in the standard-of-care regimen corresponded with an initial weight loss, highlighting the potential impact of treatment composition on both health status and tolerability.

## DISCUSSION

The current recommendation for treating *M. avium* disease is the simultaneous administration of multiple drugs—commonly clarithromycin, ethambutol, and rifampicin—which together constitute the standard-of-care regimen. This multidrug approach is designed not only to maximize efficacy but also to minimize the likelihood of resistance emergence, a major cause of refractory disease in some MAC subtypes [3,6,10]. Indeed, a recent clinical study evaluating patients who received more than 12 months of therapy showed that the combination of clarithromycin, ethambutol, and rifampicin was associated with higher odds of microbiological cure compared to maintenance with clarithromycin alone [35]. Clinical experiences and findings such as this reinforce the rationale for combining drugs in the treatment of *M. avium* disease. However, it remains uncertain whether non-inferior efficacy— without compromising the risk for resistance emergence—can be achieved using regimens that involve fewer drugs at any one time. Miwa and colleagues demonstrated noninferiority of clarithromycin+ethambutol dual therapy compared to clarithromycin+ethambutol+rifampicin in select populations with reduction in adverse events and treatment discontinuation in the dual-therapy group, but data on resistance development or relapse were not reported [36]. Such approaches could reduce the cumulative risk of drug-associated adverse events, which are a leading cause of treatment discontinuation and non-compliance in patients with nontuberculous mycobacterial infections [3]. Intermittent macrolide therapy has been a mainstay of noncavitary MAC treatment when used as part of a multidrug regimen, improving tolerability without reduction in sputum culture conversion rates [6,10] and implying that other closely-pulsed regimens may also avoid the specter of resistance. A sequencing-based treatment strategy, in which individual drugs are rotated rather than administered concurrently, could theoretically balance efficacy with improved tolerability, but this has not been adequately tested.

Our study serves as proof-of-concept for sequenced monotherapy of *M. avium* disease. It is generally accepted that monotherapy increases the likelihood of selecting resistant subpopulations, since spontaneous resistant mutants—though rare at the outset—can expand when drug pressure eliminates susceptible bacteria [37]. By the conclusion of monotherapy, this shift in population dynamics typically enables easier detection of resistant clones. In our study, however, random sampling of two colonies per mouse at each timepoint, followed by MIC determination for all drugs tested, did not reveal mutants that could be classified as resistant. This suggests that sequenced monotherapy, in contrast to continuous monotherapy, does not carry a high risk of resistance selection over a three-month period. Although our study was not designed to test single-agent therapy in isolation, these findings provide preliminary evidence that sequencing drugs may mitigate the resistance risks usually attributed to monotherapy over extended periods.

There are, nonetheless, important limitations in our study. First, it is important to note the exclusion of cavitary disease at treatment initiation. Pre-existing cavitation would be expected to dramatically alter drug delivery to select mycobacterial subpopulations and promote the development of resistance to any single- or multi-agent regimen. This multifactorial process is difficult to model in mice, is not observed in this model, and was beyond the scope of our current focus on non-complex, treatment-naïve *M. avium* pulmonary disease. As a proof-of-concept investigation, we restricted our design to three drugs—clarithromycin, bedaquiline, and clofazimine—although current guidelines recommend additional agents [6,10]. There is emerging interest in developing regimens with bedaquiline and clofazimine to treat *M. avium* disease [21,38].

With three drugs, six unique monotherapy sequences are theoretically possible, yet we tested only two: clarithromycin → bedaquiline → clofazimine and the reverse. Both yielded comparable efficacy and MIC outcomes, suggesting that other sequences may not fundamentally alter results. However, whether starting or ending with bedaquiline, given its unique mechanism and sterilizing activity, might produce distinct outcomes warrants further investigation. Moreover, because both bedaquiline and clofazimine have plasma half-lives lasting several weeks, their persistence after dosing effectively extends multidrug exposure beyond the planned treatment period. Additionally, our study did not include sequenced monotherapy evaluations of the drugs used in the standard-of-care regimen.

In our previous study conducted under identical experimental conditions, we evaluated the efficacy of the same three drugs used here—clarithromycin, bedaquiline, and clofazimine—administered as combination and as monotherapies at the same doses for eight weeks [21]. During the first month, the triple-drug combination (clarithromycin+bedaquiline+clofazimine) achieved the greatest reduction in *M. avium* burden. However, at the conclusion of eight weeks of treatment, the dual regimen (clarithromycin+bedaquiline) produced a reduction in lung CFU comparable to the triple-drug combination. As controls, clarithromycin and bedaquiline monotherapies were also evaluated; although less potent, both continued to lower *M. avium* burden throughout the study. These findings motivated our current investigation into the efficacy of sequenced monotherapy.

Next, our resistance assessment relied on sampling 10 colonies from each treatment group at the end of each phase. While this represents only ∼0.1% of colonies at the first four-week timepoint, the proportion increased to ∼10% and ∼40–60% at the subsequent two timepoints, making sampling more representative later in therapy (Figure 1). Although the absence of MIC increases supports the conclusion that resistance did not emerge, larger-scale sampling at the conclusion of the first four weeks of treatment would strengthen confidence in this observation for this timepoint. A potential concern could be that drugs with long half-lives might interfere with CFU enumeration by inhibiting bacterial growth in agar after plating lung homogenates. To minimize this carryover effect, we plated 10-fold serial dilutions of the lung homogenates, thereby greatly reducing residual drug concentrations in the samples used for *M. avium* CFU quantification. Finally, broader exploration of sequenced monotherapy should include additional drugs, variations in dose and frequency, different monotherapy durations, and diverse MAC clinical isolates. Such refinements, while beyond the scope of the first proof-of-concept study such as this, will be critical to evaluate generalizability and clinical feasibility.

In summary, our findings provide initial evidence that sequenced monotherapy in a mouse model of chronic *M. avium* lung infection can preserve efficacy comparable to standard multidrug therapy without increasing the risk of resistance emergence. Importantly, our results do not challenge the well-established view that continuous monotherapy promotes resistance; rather, they highlight that sequenced monotherapy represents a distinct strategy with a different risk profile. An additional potential advantage of this approach is that, by limiting exposure to only one drug at a time, sequenced monotherapy may reduce cumulative drug-related toxicities and thereby improve patient experience and treatment adherence compared to current standard-of-care regimens. As a proof-of-concept study, this work establishes a foundation for more comprehensive investigations designed to test the robustness of this approach. If further testing with drugs not included in this study demonstrates that sequenced monotherapy achieves non-inferior efficacy, a clinical trial would be warranted to evaluate the clinical translatability of this approach.

## Author contributions

RAH: investigation, data analysis and interpretation, manuscript preparation. BR: investigation, data analysis and interpretation, manuscript preparation. JK: methodology and data analysis. CMP: methodology and investigation. GL: study conception, study design, project administration, data interpretation, manuscript preparation, and funding acquisition.

## Disclaimer

The National Institutes of Health (NIH) and Sherrilyn and Ken Fisher Center for Environmental Infectious Diseases played no role in the design and conduct of the study; generation, analysis and interpretation of data; and in the preparation and approval of the manuscript. The content is solely the responsibility of the authors.

## SUPPLEMENTARY DATA

## SUPPLEMENTARY TABLE

**Supplementary Table 1:**
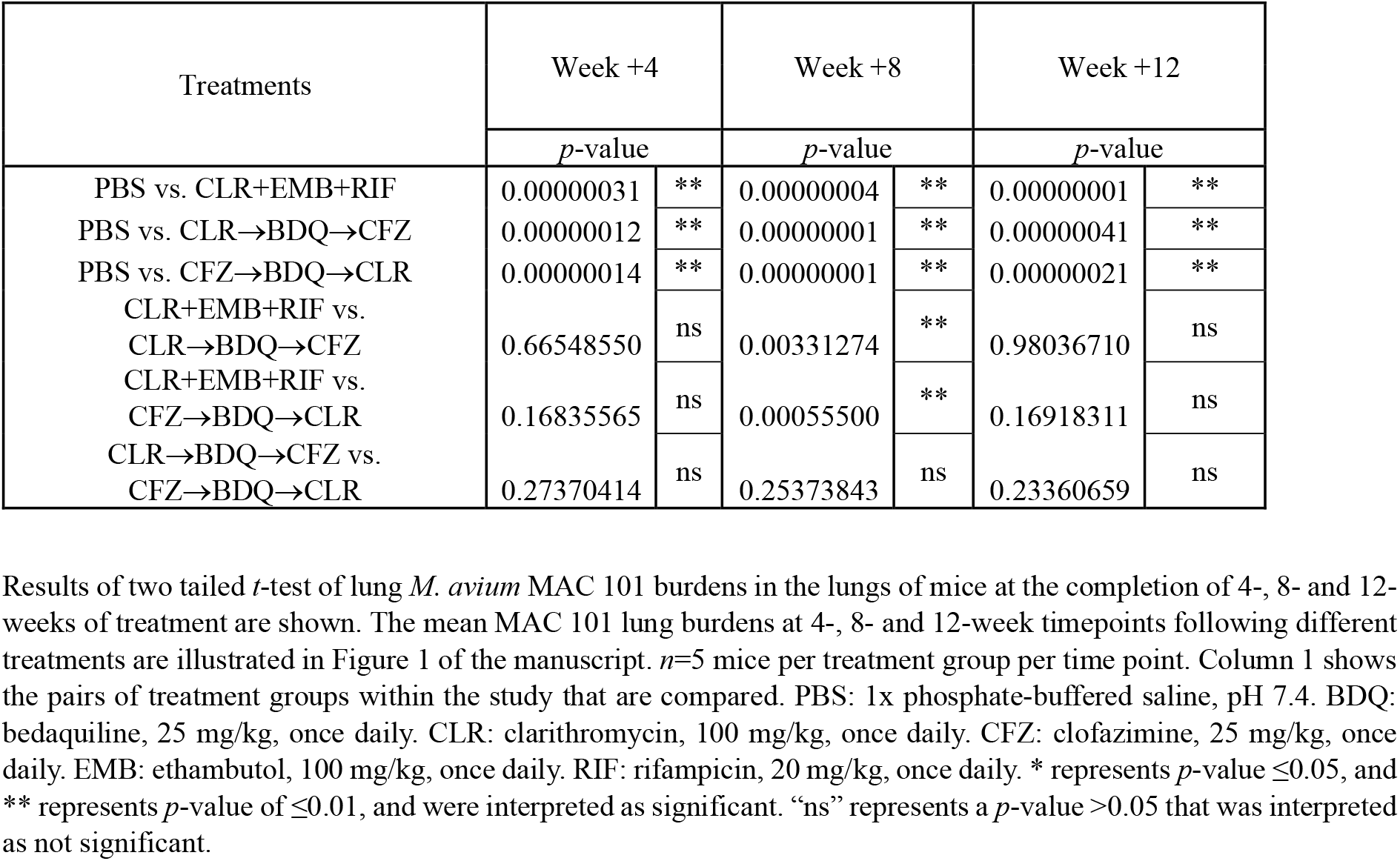
Statistical assessment of lung *M. avium* burden between groups of mice receiving different treatments shown in figure 1.

| Treatments | Week +4 |  | Week +8 |  | Week +12 |  |
| --- | --- | --- | --- | --- | --- | --- |
|  | *p*-value |  | *p*-value |  | *p*-value |  |
| PBS vs. CLR+EMB+RIF | 0.00000031 | ** | 0.00000004 | ** | 0.00000001 | ** |
| PBS vs. CLR→BDQ→CFZ | 0.00000012 | ** | 0.00000001 | ** | 0.00000041 | ** |
| PBS vs. CFZ→BDQ→CLR | 0.00000014 | ** | 0.00000001 | ** | 0.00000021 | ** |
| CLR+EMB+RIF vs.<br>CLR→BDQ→CFZ | 0.66548550 | ns | 0.00331274 | ** | 0.98036710 | ns |
| CLR+EMB+RIF vs.<br>CFZ→BDQ→CLR | 0.16835565 | ns | 0.00055500 | ** | 0.16918311 | ns |
| CLR→BDQ→CFZ vs.<br>CFZ→BDQ→CLR | 0.27370414 | ns | 0.25373843 | ns | 0.23360659 | ns |

**Supplementary Table 2:** Non-inferiority test comparing experimental treatments with the standard-of-care.

| CLR→BDQ→CFZ vs CLR+EMB+RIF |  |  | CFZ→BDQ→CLR vs CLR+EMB+RIF |  |
| --- | --- | --- | --- | --- |
| Mean CFU-experimental treatment | 1.8 |  | Mean CFU-experimental treatment | 1.56 |
| Mean CFU-standard-of-care | 1.8 |  | Mean CFU-standard-of-care | 1.80 |
| Standard deviation-experimental treatment | 0.31 |  | Standard deviation-experimental treatment | 0.3 |
| Standard deviation- standard-of-care | 0.2 |  | Standard deviation- standard-of-care | 0.2 |
| N-experimental treatment | 5 |  | N-experimental treatment | 5 |
| N- standard-of-care | 5 |  | N- standard-of-care | 5 |
| Non-inferiority margin ($\Delta$) | -0.3 | | Non-inferiority margin ($\Delta$) | -0.3 |
| Alpha (one-sided) | 0.05 |  | Alpha (one-sided) | 0.05 |
| Difference in means | 0 |  | Difference in means | 0.24 |
| Standard error | 0.165 |  | Standard error | 0.161 |
| Z for 95% CI (one-sided) | 1.645 |  | Z for 95% CI (one-sided) | 1.645 |
| Lower bound of CI | -0.271 |  | Lower bound of CI | -0.025 |
| **Conclusion: Non-inferior** |  |  | **Conclusion: Non-inferior** |  |

Result of non-inferiority test with 15% margin comparing lung *M. avium* colony forming unit burdens at the conclusion of 12 weeks of experimental treatments and standard-of-care. Experimental treatments include sequenced monotherapies of clarithromycin monotherapy for four weeks followed by bedaquiline monotherapy for four weeks, followed by clofazimine monotherapy for the final four weeks (CLR→BDQ→CFZ) and the same drugs and doses but administered in reverse order (CFZ→BDQ→CLR). The standard-of-care was a triple-drug combination comprising clarithromycin+ethambutol+rifampicin (CLR+EMB+RIF).

**Supplementary Table 3:** Statistical assessment of mean body weights between groups of mice receiving different treatments shown in figure 3.

| Treatments | Week +2 |  | Week +4 |  | Week +6 |  | Week +8 |  | Week +10 |  | Week +12 |  |
| --- | --- | --- | --- | --- | --- | --- | --- | --- | --- | --- | --- | --- |
|  | *p*-value |  | *p*-value |  | *p*-value |  | *p*-value |  | *p*-value |  | *p*-value |  |
| PBS vs. CLR+EMB+RIF | 0.206 | ns | 0.811 | ns | 0.218 | ns | 0.858 | ns | 0.215 | ns | 0.464 | ns |
| PBS vs. CLR+BDQ+CFZ | 0.082 | ns | 0.099 | ns | 0.776 | ns | 0.647 | ns | 0.620 | ns | 0.033 | * |
| PBS vs. CFZ+BDQ+CLR | 0.618 | ns | 0.811 | ns | 0.546 | ns | 0.603 | ns | 0.558 | ns | 0.757 | ns |
| CLR+EMB+RIF vs. CLR+BDQ+CFZ | 0.045 | * | 0.101 | ns | 0.282 | ns | 0.711 | ns | 0.168 | ns | 0.013 | * |
| CLR+EMB+RIF vs. CFZ+BDQ+CLR | 0.300 | ns | 0.726 | ns | 0.779 | ns | 0.569 | ns | 0.199 | ns | 0.918 | ns |
| CLR+BDQ+CFZ vs. CFZ+BDQ+CLR | 0.044 | * | 0.245 | ns | 0.498 | ns | 0.468 | ns | 0.837 | ns | 0.121 | ns |

